# CLASPs stabilize the intermediate state between microtubule growth and catastrophe

**DOI:** 10.1101/2022.12.03.518990

**Authors:** EJ Lawrence, S Chatterjee, M Zanic

## Abstract

CLASPs regulate microtubules in many fundamental cellular processes. CLASPs stabilize dynamic microtubules by suppressing catastrophe and promoting rescue, the switch-like transitions between microtubule growth and shrinkage. However, the molecular mechanisms underlying CLASP’s activity are not understood. Here, we investigate the effects of CLASPs on distinct microtubule substrates in the absence of tubulin to gain insight into how CLASPs regulate microtubule dynamics. Surprisingly, we find that human CLASP1 depolymerizes stable microtubules in the presence of GTP, but not in the absence of nucleotide. Conversely, CLASP1 stabilizes dynamic microtubules upon tubulin dilution in the presence of GTP. Our results demonstrate that CLASP1 drives microtubule substrates with different inherent stabilities into the same slowly-depolymerizing state in the absence of tubulin in a nucleotide-dependent manner. We interpret this state as the pre-catastrophe intermediate state between microtubule growth and shrinkage. Thus, we conclude that CLASPs stabilize the intermediate state between microtubule growth and shrinkage to suppress microtubule catastrophe and promote rescue.

## INTRODUCTION

Microtubules are dynamic cytoskeletal polymers essential for fundamental cellular processes. Individual microtubules switch between phases of growth and shrinkage through a behavior known as microtubule dynamic instability (Mitchison and Kirschner 1984). The transition from microtubule growth to shrinkage is called catastrophe, and the transition from shrinkage back to growth is called rescue. To facilitate their roles in diverse cellular processes, microtubules are regulated by a large number of microtubule-associated proteins (MAPs). CLASPs (cytoplasmic linker-associated proteins) are a highly conserved family of microtubule-associated proteins that stabilize microtubules in many cellular contexts including cell division, cell migration and neuronal development (Akhmanova et al. 2001, Lawrence et al. 2020). Human CLASPs stabilize microtubules by autonomously suppressing microtubule catastrophe and promoting microtubule rescue without changing the rates of microtubule growth or shrinkage (Aher et al. 2018, Lawrence et al. 2018, Lawrence and Zanic 2019). However, the molecular mechanisms underlying CLASP’s activity remain largely unknown.

CLASPs belong to a larger group of proteins that use tubulin-binding TOG (tumor overexpression gene) domains to modulate microtubule dynamics (Slep 2009, Al-Bassam and Chang 2011, Farmer and Zanic 2021). Another prominent TOG-domain protein, XMAP215, uses its TOG domains to accelerate microtubule growth (Gard and Kirschner 1987, Brouhard et al. 2008). XMAP215’s mechanism involves stabilizing a weakly-bound tubulin dimer at the microtubule end, which serves as an intermediate state in microtubule polymerization (Ayaz et al. 2012, Ayaz et al. 2014, Brouhard and Rice 2014). In contrast, human CLASPs do not affect the microtubule growth rate, but specifically modulate microtubule catastrophe and rescue (Aher et al. 2018, Lawrence et al. 2018, Lawrence and Zanic 2019). Additionally, the unique architecture of CLASPs TOG domains suggests that CLASPs regulate a distinct tubulin conformation (Leano et al. 2013, Maki et al. 2015, Leano and Slep 2019, Lawrence et al. 2020). It is not known what state of the microtubule end CLASPs act on to regulate the transitions between microtubule growth and shrinkage.

Interestingly, an early model of microtubule dynamic instability proposed the existence of a metastable intermediate state of dynamic instability between the microtubule growth and shrinkage phases (Tran et al. 1997, Janosi et al. 2002). In support of this, recent work found that dynamically growing microtubules exhibit a distinct slowdown in growth prior to catastrophe (Maurer et al. 2014, Mahserejian et al. 2022). Importantly, in the presence of CLASP, microtubules can withstand large growth fluctuations, and return to a robust growth phase following transient growth slowdowns, thus avoiding catastrophe (Lawrence et al. 2018, Mahserejian et al. 2022). Furthermore, CLASPs promote microtubule pausing in cells and *in vitro*, and stabilize microtubule ends at anchor points such as kinetochores, focal adhesions, and the cell cortex (Sousa et al. 2007, Li et al. 2012, Aher et al. 2018, Lawrence et al. 2020, Mahserejian et al. 2022). Therefore, we hypothesized that CLASPs stabilize microtubules in the pre-catastrophe state, an intermediate state between microtubule growth and shrinkage.

The stabilization of an intermediate state drives a biochemical reaction in either forward or reverse direction, depending on the availability of the reactants. For XMAP215, the absence of soluble tubulin reverses its activity from a microtubule polymerase to depolymerase (Shirasu-Hiza et al. 2003, Brouhard et al. 2008). This discovery provided critical insight into XMAP215’s mechanism of action. To what extent CLASP’s anti-catastrophe activity may be modulated by the availability of soluble tubulin is not known. Here, we investigate the effects of CLASPs on distinct microtubule substrates in the absence of tubulin to unravel the molecular mechanisms by which CLASPs regulate microtubule catastrophe and rescue.

## RESULTS

### Human CLASP1 depolymerizes GMPCPP-stabilized microtubules in a GTP-dependent manner

To investigate whether CLASP1, like XMAP215, has the ability to depolymerize stabilized microtubules, we used an established *in vitro* assay, combining TIRF microscopy with purified protein components (Gell et al. 2010). Briefly, stable microtubules were polymerized with GMPCPP, a slowly hydrolyzable GTP analogue, and adhered to coverslips. Microtubule depolymerization was monitored over 15 minutes under different reaction conditions (Figure 1A). In the buffer control condition, in the absence of any MAPs, microtubules depolymerized very slowly over the course of the experiment, as expected (Figure 1B left, 1D; 0.29 nm/s ± 0.04 nm/s; SE, N = 90). As a positive control, we purified recombinant chTOG (Figure S1), the human homolog of XMAP215, and tested its depolymerase activity. Indeed, we observed microtubule depolymerization with 200 nM chTOG (Figure 1B middle; 2.64 nm/s ± 0.02 nm/s; SE, N = 50). Thus, like XMAP215, human chTOG depolymerizes microtubules in the absence of soluble tubulin. In contrast, when stabilized microtubules were incubated with 200 nM purified recombinant CLASP1 (Figure S1), we did not observe significant microtubule depolymerization (Figure 1B right; 0.248 nm/s ± 0.008 nm/s; SE, N = 79; p = 0.59 compared to the buffer control). Thus, unlike XMAP215/chTOG, CLASP1 does not depolymerize microtubules under these conditions.

**Figure 1.**
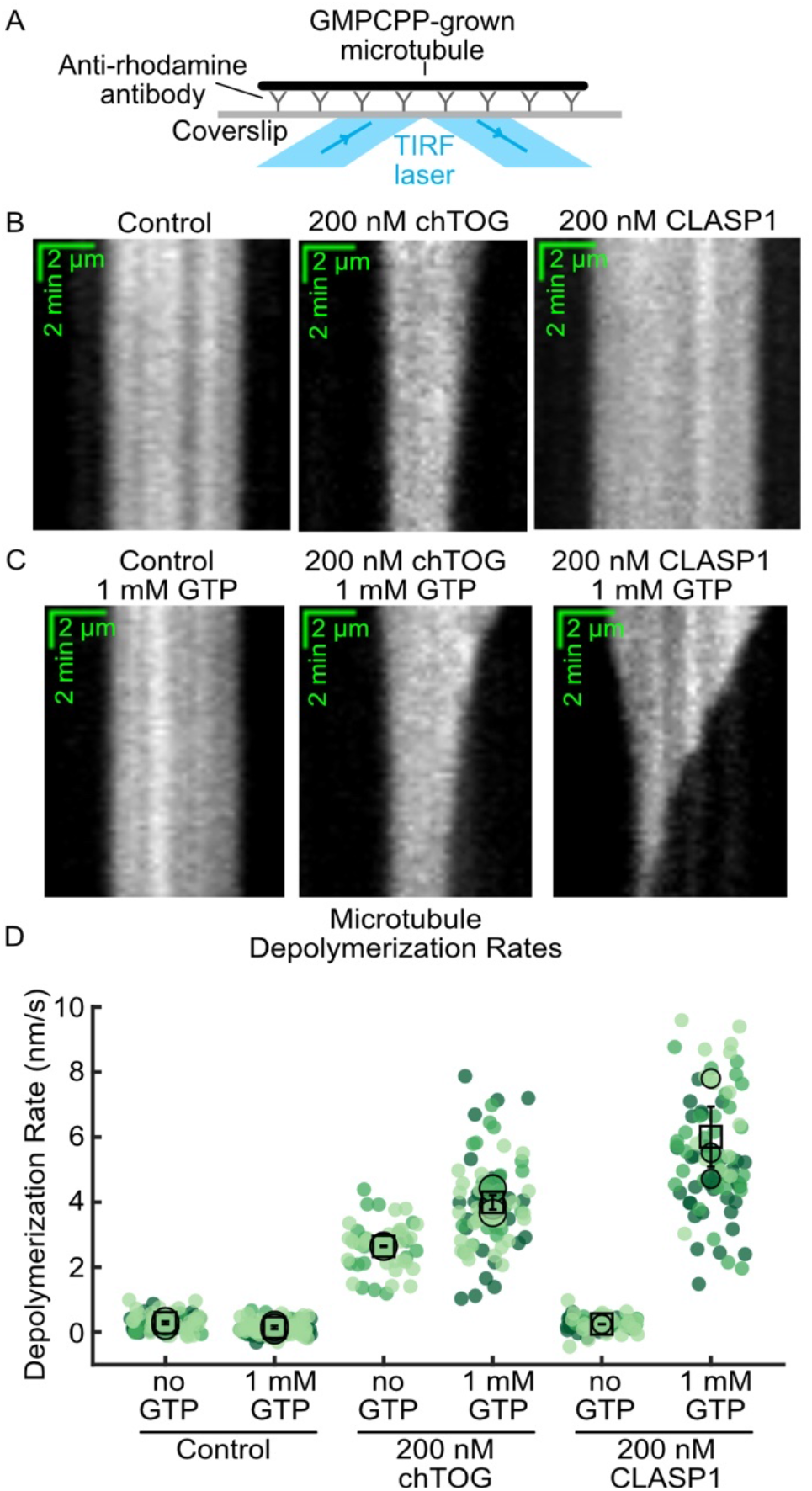
Human CLASP1 depolymerizes GMPCPP-stabilized microtubules in a GTP-dependent manner. (A) Schematic of the microtubule depolymerization assay. (B) Representative kymographs of GMPCPP-stabilized microtubules incubated with storage buffer, 200 nM chTOG or 200 nM CLASP1. (C) Representative kymographs of GMPCPP-stabilized microtubules incubated with storage buffer, 200 nM chTOG, or 200 nM CLASP1 in the presence of 1 mM GTP. (D) Quantification of microtubule depolymerization rates for the conditions in (B) and (C). N = 50 - 108 microtubules for each condition across at least 2 experimental days. Individual data points from different experiments are plotted in different shades and the means for each experimental repeat are plotted as larger points in the same color. The squares indicate the average of the experimental means and the vertical bars are the standard errors of the means.

The configuration of microtubule ends depends on the nucleotide state of the end-bound tubulin dimers. Given that the specific configuration of tubulin recognized by CLASP’s TOG domains is not known, we wondered whether CLASP1’s activity is sensitive to the nucleotide state of the tubulin at the microtubule ends. Therefore, we investigated the effects of including GTP in the reaction mix for our depolymerization assay, which is expected to exchange with the GMPCPP bound to the terminal tubulin dimers (Mitchison 1993) (Figure 1B). The addition of 1 mM GTP to the buffer control condition did not promote microtubule depolymerization (Figure 1B left, 1C; 0.14 nm/s ± 0.05 nm/s; SE, N = 108; p = 0.25 compared to the no GTP control). Therefore, the presence of GTP itself does not significantly destabilize the microtubule lattice. Furthermore, chTOG displayed only a mild, 1.5-fold increase in depolymerase activity in the presence of 1 mM GTP compared to the no GTP condition (Figure 1B middle, 1C; 4.0 nm/s ± 0.2 nm/s; SE, N = 77). In contrast, upon introduction of 200 nM CLASP1 and 1 mM GTP, the microtubules robustly depolymerized, displaying a 24-fold increase in depolymerization rate when compared to the CLASP1 with no GTP condition (Figure 1B right, Figure 1C; Supplementary Movie 1; 6.0 nm/s ± 0.9 nm/s; SE, N = 89; p = 0.024 compared to CLASP1 without GTP). Overall, we conclude that similar to XMAP215 and chTOG, CLASP1 robustly depolymerizes stabilized microtubules in the absence of soluble tubulin. However, while chTOG’s depolymerase activity shows little sensitivity to the presence of GTP in solution, CLASP1’s activity is strongly promoted by GTP. These results demonstrate that, in the absence of soluble tubulin, CLASP1 possesses GTP-dependent microtubule depolymerase activity.

### A minimal TOG2 domain construct is sufficient for microtubule depolymerase activity

Humans possess two CLASP paralogs: CLASP1 and CLASP2, with multiple TOG domains contained within all major isoforms (CLASP1α, CLASP2α, and CLASP2**γ**) (Figure 2A). Previous work established that a minimal construct composed of the TOG2 domain and the serine-arginine-rich region of CLASP2α (TOG2-S) recapitulates the anti-catastrophe and rescue activity of full-length human CLASP2α on dynamically-growing microtubules (Aher et al. 2018). We, therefore, set out to establish whether other CLASP isoforms depolymerize microtubules in the presence of GTP, and determine the minimal requirements for CLASP’s depolymerase activity. To address this, we incubated GMPCPP-stabilized microtubules with 200 nM of CLASP1, CLASP2α, or CLASP2**γ** and 1 mM GTP (Figure 2B). While CLASP1 displayed the strongest depolymerase activity, we found that all CLASP family members depolymerized the stabilized microtubules in the presence of GTP (Figure 2B-C). Next, we tested whether the isolated TOG2-S construct also possesses microtubule depolymerase activity. Indeed, we observed microtubule depolymerization when microtubules were incubated with 200 nM EGFP-tagged TOG2-S and 1 mM GTP (Figure 2B-C). The depolymerization rates of microtubules with all CLASP family members and the minimal TOG2-S domain construct were statistically significantly different from the control (p<0.001, one-way ANOVA followed by a post hoc Tukey HSD). The finding that the minimal TOG2-S construct is capable of depolymerizing stable microtubules demonstrates that CLASP’s depolymerase activity does not strictly require a string of multiple TOG domains. Interestingly, the depolymerization rates with TOG2-S were significantly different from those with CLASP2α, but not CLASP2γ, suggesting that the TOG1 domain may contribute to CLASP’s depolymerizing activity. Furthermore, the fact that TOG2-S is also the minimal unit required for CLASP’s anti-catastrophe and rescue activities suggests that the molecular mechanisms underlying CLASP’s depolymerase activity are linked to its mechanism of microtubule dynamics regulation.

**Figure 2.**
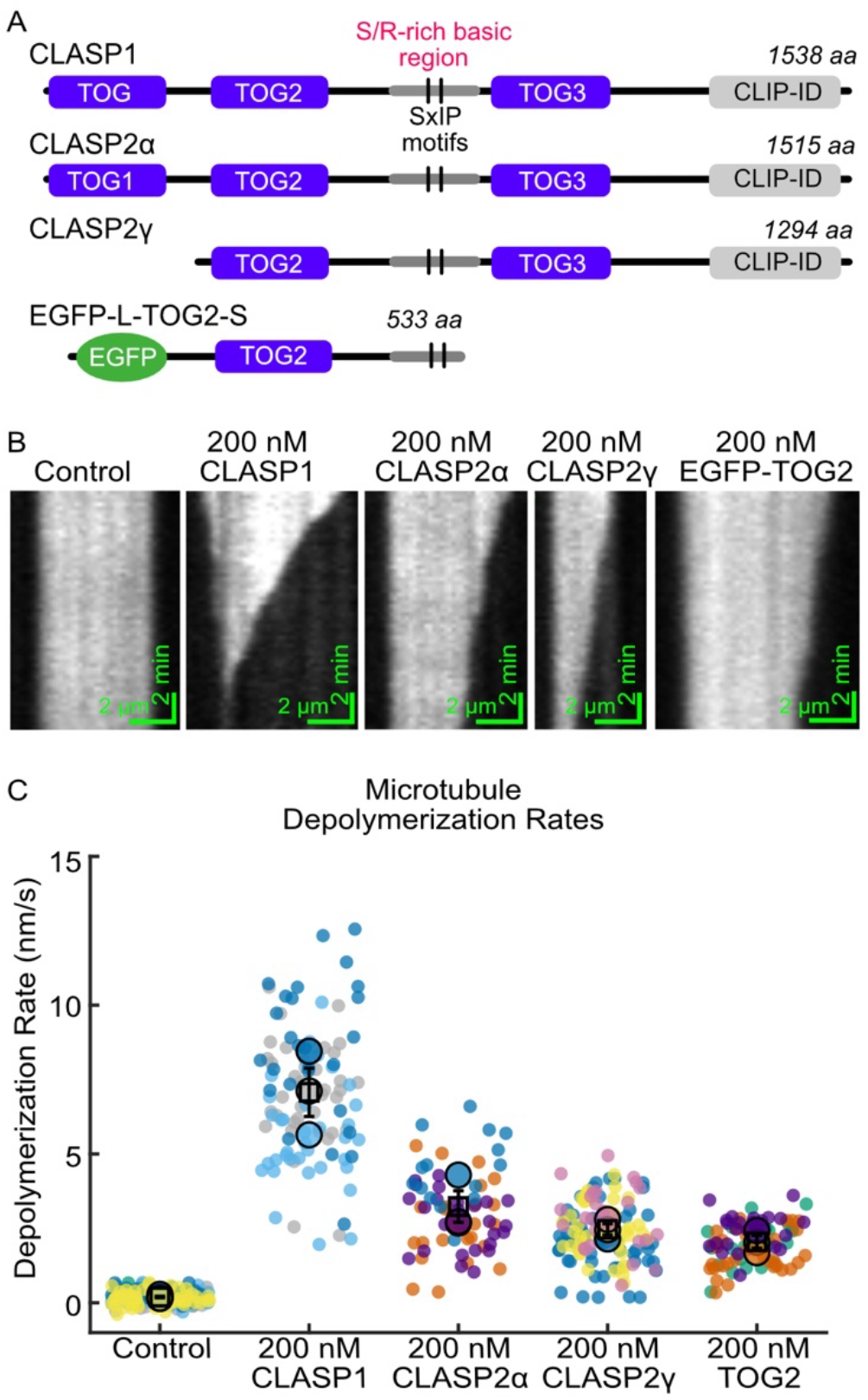
A minimal TOG2 domain construct is sufficient for microtubule depolymerase activity. (A) Domain structures of human CLASP family members and an EGFP-tagged TOG2 domain from CLASP2α. (B) Representative kymographs of GMPCPP-stabilized microtubules incubated with storage buffer, 200 nM CLASP1, 200 nM CLASP2α, 200 nM CLASP2γ, or 200 nM EGFP-L-TOG2-S in the presence of 1 mM GTP. (C) Quantification of microtubule depolymerization rates for the conditions in (B). The mean rates of microtubule depolymerization are: 0.19 nm/s +/- 0.03 nm/s (SE; N=221) for the buffer control, 7.1 nm/s +/- 0.8 nm/s (SE; N=102) for CLASP1, 3.2 nm/s +/- 0.5 nm/s (SE; N=70) for CLASP2α, 2.4 nm/s +/- 0.2 nm/s (SE; N=136) for CLASP2γ, and 2.0 +/- 0.2 nm/s (SE; N=86) for TOG2-S. All data were obtained across at least 3 different experimental days.

### CLASP1 depolymerizes plus and minus ends in the presence of GTP, but is plus-end specific in the presence of GDP

Microtubules have two structurally and biochemically distinct ends; a plus end and a minus end. In our depolymerization assays, we noticed intriguing differences in the behavior of the two microtubule ends in the presence of CLASP1 and GTP. To determine the end-specific activity of CLASP1, we performed a titration of CLASP1 concentration from 0 nM to 500 nM in the presence of 1 mM GTP on polarity-marked microtubules, allowing us to distinguish plus and minus ends (see Methods) (Figure 3A-C, Supplementary Movie 2). We found that the microtubule depolymerization rate increased with increasing CLASP1 concentrations at both plus and minus ends (Figure 3B-C). Fitting the CLASP1 titration data to the Michaelis-Menten equation revealed that the half-maximum depolymerization rate was achieved at 15 nM CLASP1 (95% CI: 10 nM – 20 nM) for plus ends, and 42 nM (95% CI: 5 nM - 79 nM) for minus ends (Figure 3C). Therefore, CLASP1 depolymerizes both microtubule plus and minus ends in the presence of GTP but the plus ends are more susceptible to CLASP1 depolymerase activity than the minus ends.

**Figure 3.**
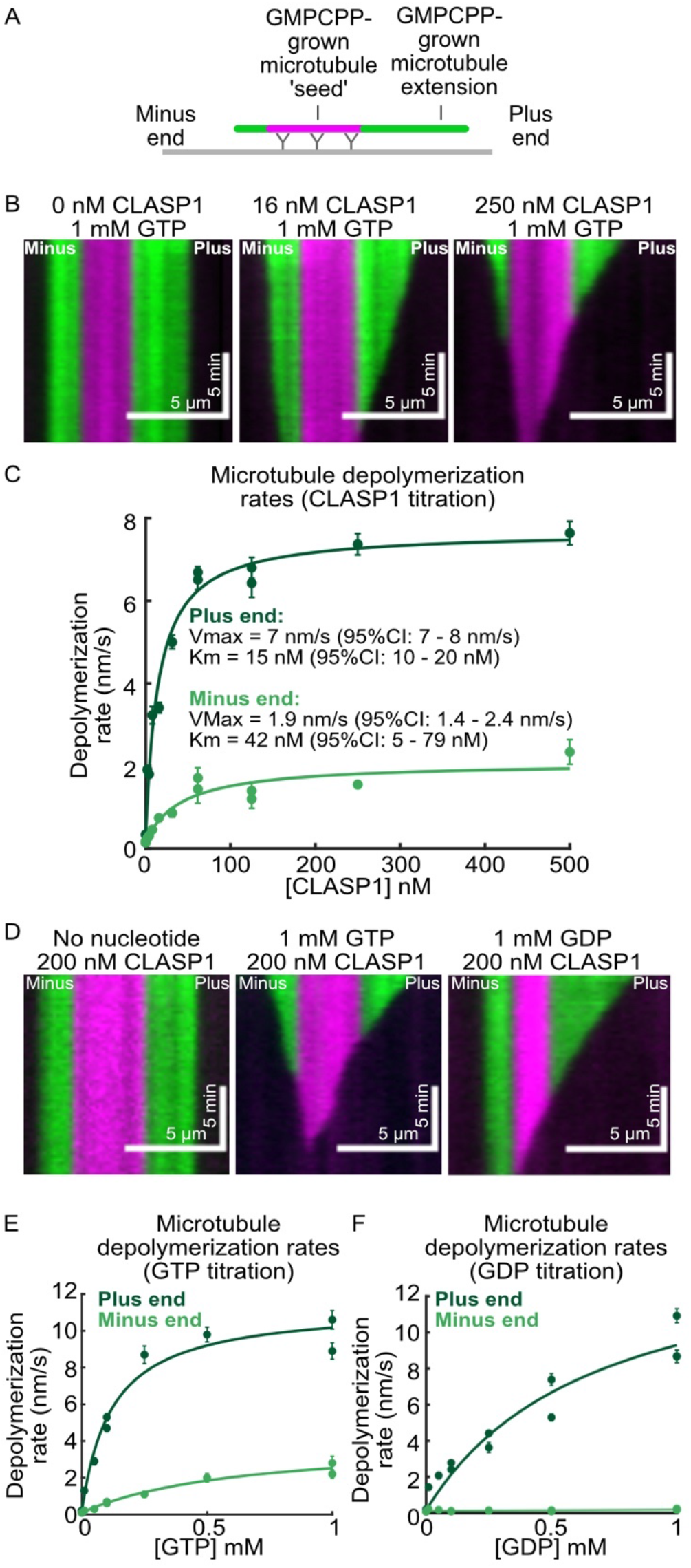
CLASP1 depolymerizes plus and minus ends in the presence of GTP, but is plus-end specific in the presence of GDP. (A) Schematic of the polarity-marked microtubule assay used to distinguish plus and minus ends. (B) Representative kymographs of polarity-marked microtubules incubated with 0 nM, 16 nM, and 250 nM CLASP1. (C) Quantification of microtubule depolymerization rates as a function of CLASP1 concentration in the presence of 1\ mM GTP. The plus end data are in dark green and the minus end data are in light green. The solid lines represent the Michaelis-Menten fit to the data. (D) Representative kymographs of polarity-marked microtubules incubated with 200 nM CLASP1 in the no nucleotide, 1 mM GTP, and 1 mM GDP conditions. (E) Quantification of microtubule depolymerization rates as a function of GTP concentration in the presence of 200 nM CLASP1. The plus end data are in dark green and the minus end data are in light green. The solid lines represent the Michaelis-Menten fit to the data. (F) Quantification of microtubule depolymerization rates as a function of GDP concentration in the presence of 200 nM CLASP1. The plus end data are in dark green and the minus end data are in light green. The solid lines represent the Michaelis-Menten fit to the data for the plus end and a linear fit for the minus end.

Intriguingly, due to the head-to-tail assembly of tubulin dimers in the microtubule lattice, microtubule minus ends do not possess an exposed nucleotide, rather it is buried in the microtubule lattice (Nogales et al. 1999). Therefore, the finding that CLASP1 depolymerizes both microtubule ends raises the question of how CLASP1 depolymerizes minus ends if CLASP1’s mechanism involves sensing or exchanging the nucleotide state of terminal tubulin dimers. One possibility is that CLASP1 facilitates nucleotide exchange all along the microtubule lattice. To investigate the potential effects of CLASP1 on lattice nucleotide exchange, we performed computational modeling in which nucleotide exchange was permitted to occur either exclusively at microtubule ends or at both ends and lattice (Figure S2). We assessed the depolymerization profiles of microtubules simulated to have mean depolymerization rates matching the experimentally observed microtubule depolymerization rates. When nucleotide exchange was simulated to occur at both ends and lattice, the characteristics of the microtubule depolymerization were very different between simulated and experimental microtubules. Specifically, the simulated microtubules displayed a highly nonlinear depolymerization profile, with depolymerization rates accelerating over time due to the exposure of lattice-exchanged dimers. In contrast, when nucleotide exchange was permitted only at microtubule ends, the depolymerization rates were constant over time, as observed in our experimental results. Therefore, the mechanism in which CLASP1 facilitates nucleotide exchange all along the microtubule lattice does not explain the experimentally-observed minus-end depolymerization. Rather, we conclude that CLASP1 depolymerizes both plus and minus ends primarily through an end-specific mechanism.

To address the specific roles of nucleotides and nucleotide hydrolysis, we next performed sequential nucleotide exchange experiments in which the CLASP1 concentration was maintained constant but the solution was exchanged for reaction mixtures containing different nucleotides (Figure S3, Supplementary Movie 3). First, we incubated microtubules in the presence of CLASP1 without any nucleotide in the solution, and saw no depolymerization, as expected. Next, we introduced 1 mM GTP in the same observation channel and observed depolymerization. Finally, we exchanged the reaction solution to include 1 mM GMPCPP and found that depolymerization quickly stopped (Figure S3). Thus, CLASP1’s depolymerase activity can be switched on with GTP and switched off with GMPCPP. We also investigated CLASP1 activity in the presence of different nucleotides and nucleotide analogs. As before, CLASP1 robustly depolymerized microtubules in the presence of 1 mM GTP. Surprisingly, CLASP1 also robustly depolymerized microtubules in the presence of 1 mM GDP. Conversely, in the presence of GMPCPP (a mimic of the GTP-state), no microtubule depolymerization was observed with CLASP1. In the presence of GTPγS (a mimic of the post-hydrolysis GDP-Pi state), a slow but significant rate of microtubule depolymerization was observed (Figure S3; 0.61 +/- 0.06 nm/s; SE, N=60; p<0.001 when compared with the GMPCPP condition). Taken together these data suggest that the post-hydrolysis state of GTP facilitates CLASP-mediated microtubule depolymerization, but that GTP hydrolysis itself is not strictly required.

Interestingly, we observed that, while CLASP1 depolymerized both of the microtubule ends in the presence of GTP, only one end appeared to depolymerize in the presence of GDP (Figure S3). To investigate this further, we assessed the depolymerase activity of 200 nM CLASP1 on polarity-marked microtubules across a range of GTP and GDP concentrations (from 0 mM – 1 mM) (Figure 3D-F). As before, CLASP1 depolymerized both microtubule plus and minus ends with GTP in solution but only plus ends with GDP (Figure 3D). The rate of CLASP1-mediated microtubule depolymerization increased with increasing GTP concentrations at both plus and minus ends (Figure 3E). The half-maximum depolymerization rate was achieved at 0.12 mM GTP (95% CI: 0.06 mM – 0.18 mM; Michaelis-Menten fit) for plus ends, and 0.6 mM GTP (95% CI: 0.2 mM – 1.1 mM) for minus ends. Notably, CLASP1 enhanced the depolymerization of plus ends to a greater extent than minus ends: CLASP1 accelerated the depolymerization rate by 55-fold at the plus ends, and by 40-fold at the minus ends when comparing the 0 mM GTP condition to the saturating GTP concentration (Vmax) for each end. In comparison, the depolymerization rate increased in a less sensitive manner with increasing GDP concentrations at microtubule plus ends, eventually reaching a similar rate to the GTP condition at the highest nucleotide concentration tested (1 mM) (Figure 3F). Strikingly, we observed no depolymerization at minus ends with CLASP1 at any GDP concentration tested (linear fit; slope: 0.07 nm s^-1^ mM^-1^ (95%CI:(−0.01, 0.15) nm s^-1^ mM^-1^). Therefore, microtubule plus ends are more sensitive to CLASP1 in the presence of both nucleotides than minus ends. Furthermore, the finding that CLASP1 depolymerizes minus ends with GTP, but not GDP, points to potentially distinct requirements for the removal of tubulin from the microtubule minus end. Overall, the distinct behaviors of the two microtubule ends are consistent with the previous report that the kinetics of the metastable intermediate state differ significantly between plus and minus ends (Tran et al. 1997).

### CLASP1 and TOG2 binding to microtubule ends is modulated by nucleotides

We next investigated the effects of the nucleotide in solution on the CLASP1’s localization on stabilized microtubules. To this end, we incubated GMPCPP-stabilized microtubules with 1 nM Alexa 488-labeled-CLASP1 in the presence of different nucleotides. When no nucleotide was present in the solution, CLASP1 displayed clear microtubule end-binding preference (Figure 4A). The observed end-binding preference of CLASP1 is similar to previous observations of full-length CLASP2γ, which preferentially associates with the plus-ends of stabilized microtubules in the absence of GTP (Lawrence et al. 2018). We also observed instances where CLASP1 localized to both microtubule ends (e.g. see Figure 4A-B, no nucleotide condition), which may explain CLASP’s ability to depolymerize microtubule plus and minus ends. Notably, CLASP1’s enhanced end-localization was lost when GTP was present in the solution (Figure 4A-B). This is consistent with a recent study demonstrating that the localization of CLASP2α on microtubule ends is lost with GTP (Luo et al. 2022). Furthermore, we observed preferential binding of CLASP1 to microtubule ends with the addition of GMPCPP, but not with the addition of GDP (Figure 4A-B). In other words, CLASP1 accumulated at the microtubule ends only in conditions that are incompatible with CLASP-mediated microtubule depolymerization (i.e., no nucleotide or GMPCPP in solution), but did not accumulate at the microtubule ends in conditions compatible with depolymerization (i.e., GTP or GDP in solution). These observations led us to hypothesize that in the presence of GTP or GDP individual CLASP molecules dissociate from microtubule ends along with the terminal tubulin dimers.

**Figure 4.**
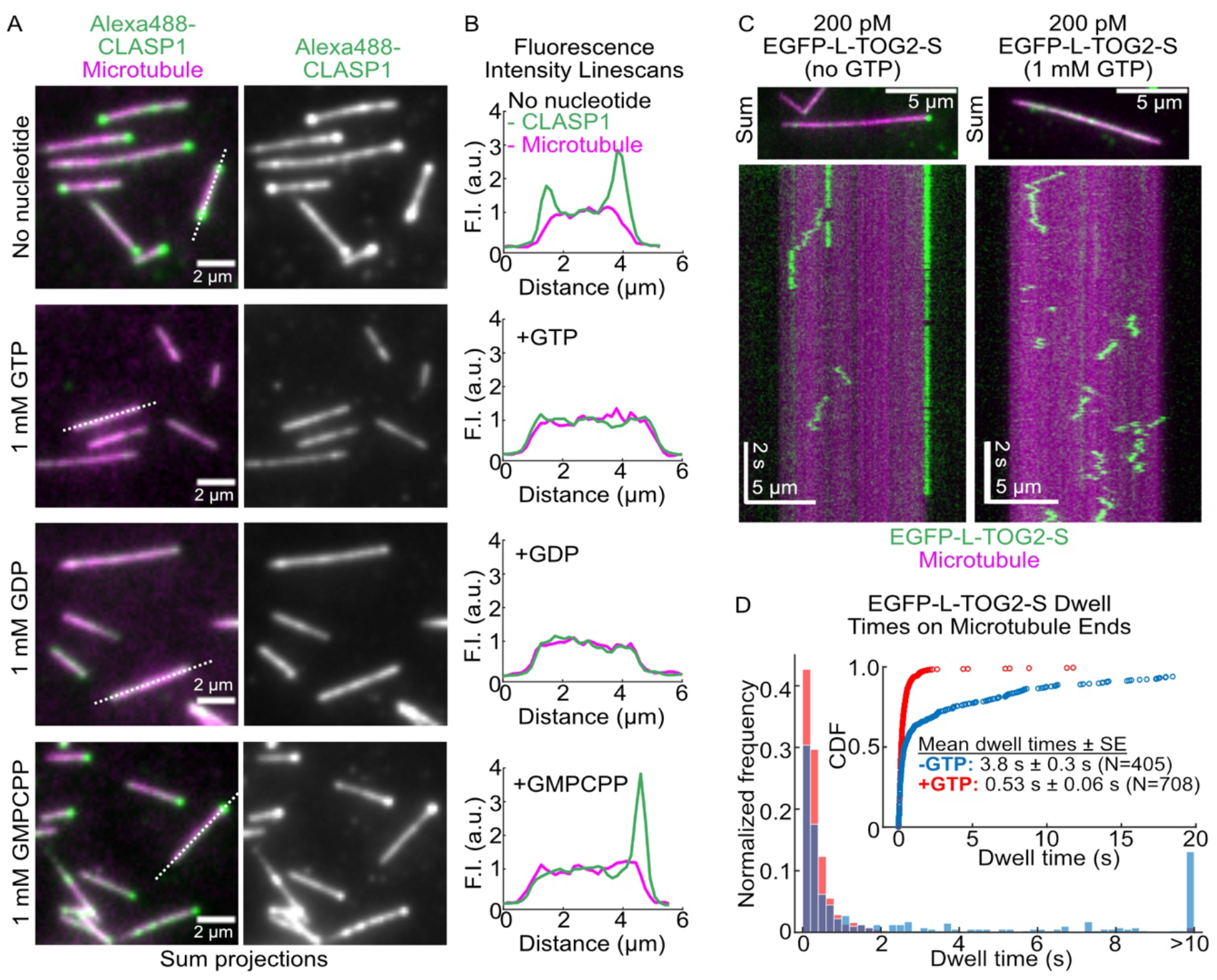
CLASP1 and TOG2 binding to microtubule ends is modulated by nucleotides. (A) Representative sum projection images of 1 nM Alexa488-CLASP1 on GMPCPP-stabilized microtubules in the absence or presence of the indicated nucleotide. Images are sum-projections of the 488-CLASP1 intensities from the first 5 seconds (100 frames) of 30-second movies imaged at 20 fps. The dotted lines indicate the positions of the corresponding linescans. (B) Fluorescence intensity linescans of the microtubules indicated by the dotted lines in (A). (C) Representative sum projection images of GMPCPP-stabilized microtubules incubated with 200 pM EGFP-L-TOG2-S with and without 1 mM GTP. The top images are sum projections from the first 5 seconds (100 frames) of 30-second movies imaged at 20 fps. Below are representative kymographs of GMPCPP-stabilized microtubules incubated with 200 pM EGFP-L-TOG2-S with and without 1 mM GTP and imaged at 20 fps. (D) Quantification of the single-molecule EGFP-L-TOG2-S dwell times on GMPCPP-stabilized microtubules in the presence and absence of 1 mM GTP. The inset shows the cumulative distribution function of the data. Data are from 3 independent experimental repeats.

To gain further insight into the depolymerase activity on a molecular level, we investigated whether GTP affects the duration of single-molecule TOG2-S binding events on microtubules. GMPCPP-stabilized microtubules were incubated with 200 pM EGFP-L-TOG2-S in the presence and absence of 1 mM GTP and imaged at 20 fps. We observed that TOG2-S also exhibited microtubule end preference in the absence of GTP (Figure 4C). In the absence of GTP, the dwell times of EGFP-L-TOG2-S were long, with a mean dwell time of 3.8 ± 0.3 s (SE, N = 405). In contrast, in the presence of GTP, the dwell times of EGFP-L-TOG2-S were significantly shorter (0.53 ± 0.06 s; SE, N = 708, p <0.001 compared to the no nucleotide condition, Mann-Whitney test) (Figure 4D). Our measurements of microtubule depolymerization rates in the presence of TOG2-S and GTP correspond to the removal of one tubulin dimer every ∼0.3 s on average (for an overall depolymerization rate of ∼2 nm/s, Figure 2C), remarkably similar to the single-molecule dwell times in the presence of GTP. We thus speculate that microtubule depolymerization occurs through TOG2-mediated dissociation of individual terminal tubulin dimers which have exchanged their nucleotide to GTP.

### CLASP1 dictates the microtubule stability in a nucleotide-dependent manner

Thus far, we have demonstrated that, in the absence of soluble tubulin, CLASP1 drives the depolymerization of stable microtubules in a nucleotide-dependent manner. However, in physiological conditions, CLASP operates along with tubulin in solution, and stabilizes microtubules by specifically suppressing microtubule catastrophe and promoting rescue (Aher et al. 2018, Lawrence et al. 2018, Lawrence and Zanic 2019). Indeed, we found that a titration of soluble tubulin from 0 μM to 8 μM in the presence of 200 nM CLASP1 and 1 mM GTP resulted in a concentration-dependent decrease in the rate of microtubule depolymerization (Figure S4). At tubulin concentrations higher than 6 μM, microtubule depolymerization was inhibited and microtubules grew extensions that did not undergo catastrophe or shrinkage during the time course of the experiment. Therefore, CLASP1 activity switches from depolymerizing, when the tubulin concentration is below the critical concentration for templated nucleation, to stabilizing when the tubulin concentration is above the critical concentration.

To determine whether CLASP’s stabilizing activity requires tubulin in solution, we performed ‘tubulin dilution’ experiments with unstable microtubule lattices. It has been previously observed that microtubules exhibit a brief phase of slow depolymerization prior to catastrophe upon tubulin dilution (Duellberg et al. 2016). This phase is thought to correspond to an intermediate state in which the microtubule end contains a mixture of both GTP and GDP-bound tubulin dimers. To mimic microtubule ends in a pre-catastrophe state, we polymerized microtubules using a mixture of GTP and GMPCPP (Figure 5A, see Methods). When soluble tubulin was subsequently diluted, the mixed-lattice microtubules slowly depolymerized, consistent with being in the pre-catastrophe state. The rates of slow depolymerization were similar regardless of the nucleotide in solution (Figures 5A and C, ‘mixed-nucleotide lattice’ control conditions). Interestingly, with the addition of 20 nM CLASP1 and either 1 mM GTP or 1 mM GDP, microtubule depolymerization rates did not accelerate, and the microtubules were maintained in a prolonged slowly depolymerizing state (Figure 5A and C). Strikingly, when CLASP1 and GMPCPP were introduced to the unstable mixed-nucleotide lattice, we found that the microtubules were completely stabilized, with a mean depolymerization rate of 0.33 nm/s ± 0.04 nm/s (SE, N=26) for the CLASP1 condition versus 16.8 nm/s ± 0.7 nm/s (SE, N=30) for GMPCPP-only condition (Figures 5A and C; p=0.002). Therefore, while GMPCPP on its own is not sufficient to stabilize unstable microtubules, CLASP1 in the presence of GMPCPP is able to completely prevent microtubule depolymerization, even in the absence of soluble tubulin. This result demonstrates that CLASP’s stabilizing activity does not require soluble tubulin but does depend on nucleotide. We conclude that CLASP in the presence of GTP and GDP allows the microtubule end to persist in the intermediate state. In contrast, CLASP-mediated stabilization of microtubules in the presence of GMPCPP represents a transition from the intermediate, pre-catastrophe state, to a growth-competent microtubule state.

**Figure 5.**
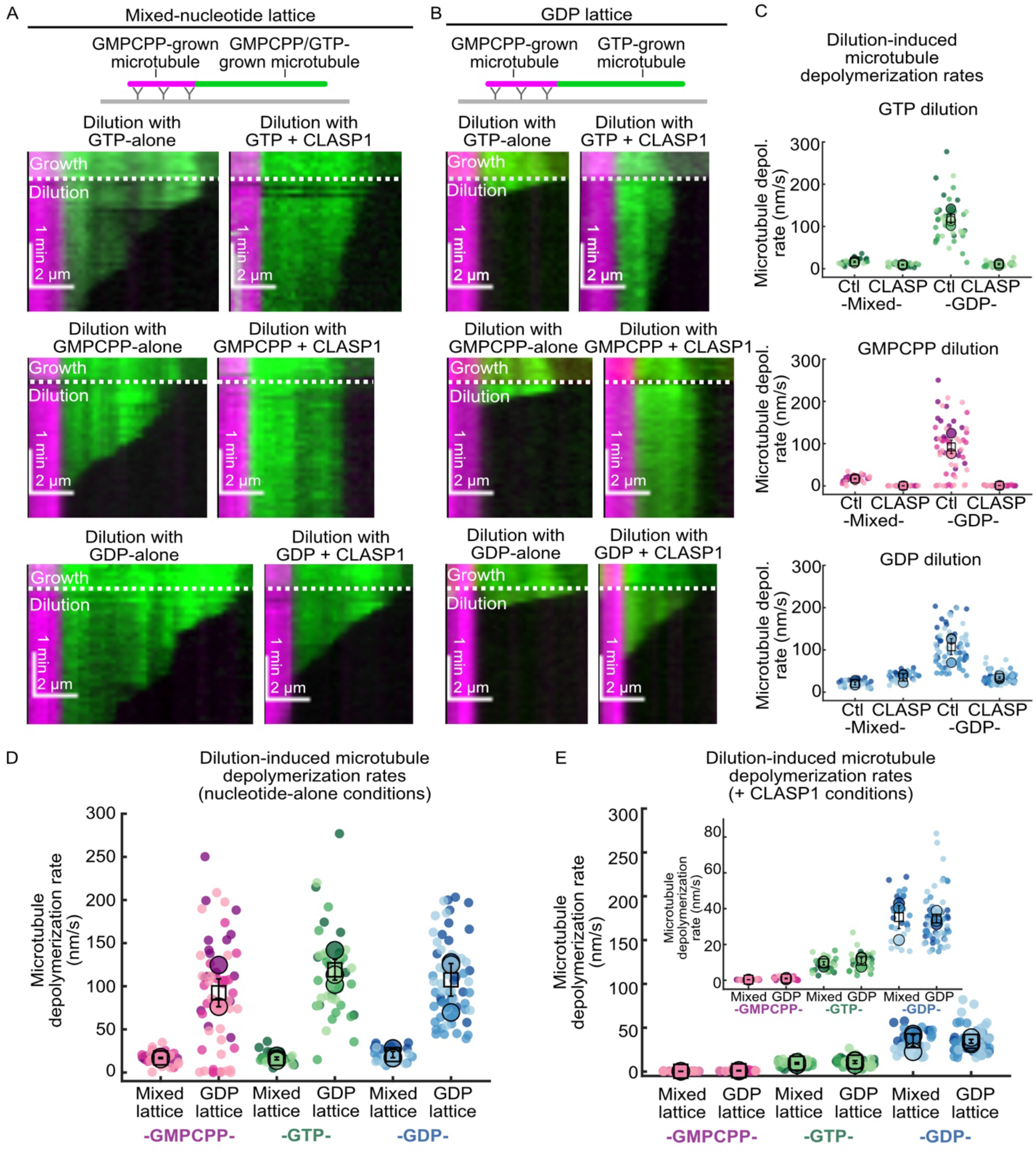
CLASP1 dictates the microtubule depolymerization rate in a nucleotide-dependent manner. (A) Representative kymographs of mixed-nucleotide microtubule extensions (green) undergoing depolymerization after tubulin dilution. Microtubules were grown with 7 μM A488 tubulin, 0.8 mM GMPCPP and 0.2 mM GTP, and then the buffer was exchanged while imaging (dotted white line) for nucleotides alone or in the presence of 20 nM CLASP1. (B) Representative kymographs of GDP microtubule extensions (green) undergoing depolymerization after tubulin dilution. Microtubules were first grown with 12 μM A488 tubulin and 100 μM GTP, and then the buffer was exchanged while imaging (dotted white line) for nucleotides alone or in the presence of 20 nM CLASP1. (C) Quantification of the mixed-lattice and GDP-lattice microtubule depolymerization rates following dilution with and without CLASP1 in the presence of different nucleotides. Data were obtained over at least 3 different experimental days. The mean rates and statistics can be found in Supplemental Table 1. The data in (C) were replotted to show dilution-induced microtubule depolymerization rates in the presence of nucleotides alone (D) and in the presence of CLASP (E) for the different microtubule templates.

To capture dynamic microtubules in the pre-catastrophe state and directly probe the effects of CLASP1, we performed tubulin dilution experiments on dynamic microtubules polymerized with GTP. Since microtubule polymerization triggers GTP-hydrolysis within the polymer, these polymerization conditions result in mostly GDP-containing, highly unstable lattices. We then exchanged the polymerization mixture for a reaction solution containing different nucleotides, alone or with CLASP1 (Figure 5B). Upon dilution of soluble tubulin with any of the nucleotides alone, the microtubules underwent catastrophe within a few seconds, followed by the onset of fast depolymerization, as expected (Figure 5B-C, ‘GDP-lattice’ control conditions). In contrast, upon dilution with CLASP1 and GTP, the microtubules were captured in the slow shrinkage phase, depolymerizing an order of magnitude slower than the control (Figures 5B and C). Dilution with CLASP1 and GDP resulted in more moderate microtubule stabilization. Furthermore, when the growth mixture was exchanged for CLASP1 and GMPCPP, the microtubules did not undergo catastrophe at all and remained stable.

Surprisingly, we noticed that the depolymerization rates of the different microtubule substrates were remarkably similar in the presence of CLASP1, despite their very different inherent stabilities in the absence of CLASP1 (Figure 5G-H). Notably, in the presence of CLASP1 and GTP, the mean depolymerization rates were 9 nm/s ± 1 nm/s (SE, N=30) for mixed-lattice microtubules, and 11 nm/s ± 2 nm/s (SE, N=33) for GDP-lattice microtubules, and were not statistically significantly different (p=0.614). Furthermore, CLASP1 also depolymerized GMPCPP-stabilized microtubules at a similar rate of 9.7 nm/s ± 0.3 nm/s (SE; N=20) (Figure 3E, plus-end data, 1 mM GTP, 200 nM CLASP1 condition). Similarly, the depolymerization rates in the presence of CLASP1 and GDP were the same on mixed-nucleotide lattices and GDP-lattices (∼ 10 nm/s; p=0.931). Finally, all tested microtubule substrates were stable in the presence of CLASP1 and GMPCPP (∼0.5 nm/s; p=0.138). Taken together, these data indicate that CLASP1 dictates the stability of microtubule ends in a pre-catastrophe state in a nucleotide-dependent manner.

## DISCUSSION

We discovered that CLASPs specifically stabilize an intermediate state of the microtubule as it transitions from growth to shrinkage. Our results show that CLASP depolymerizes stable microtubules in a nucleotide-dependent manner. On the other hand, CLASP stabilizes unstable microtubule ends even in the absence of soluble tubulin. Remarkably, we find that CLASP drives all microtubule substrates to the same slowly-depolymerizing state in the presence of GTP. We interpret this state as an intermediate state between microtubule growth and catastrophe.

CLASP’s ability to suppress the catastrophe of dynamic microtubules upon tubulin dilution demonstrates that CLASP’s anti-catastrophe mechanism does not require soluble tubulin. This is in contrast to XMAP215, which switches between polymerase and depolymerase activities depending on the availability of soluble tubulin. Furthermore, the strong nucleotide sensitivity of CLASP1 also points to key differences in their mechanisms. Taken together, our results support a mechanism whereby CLASP controls microtubule dynamics by stabilizing a metastable, nucleotide-dependent intermediate state of the microtubule end, which occurs as the microtubule transitions from growth to catastrophe (Figure 6).

**Figure 6.**
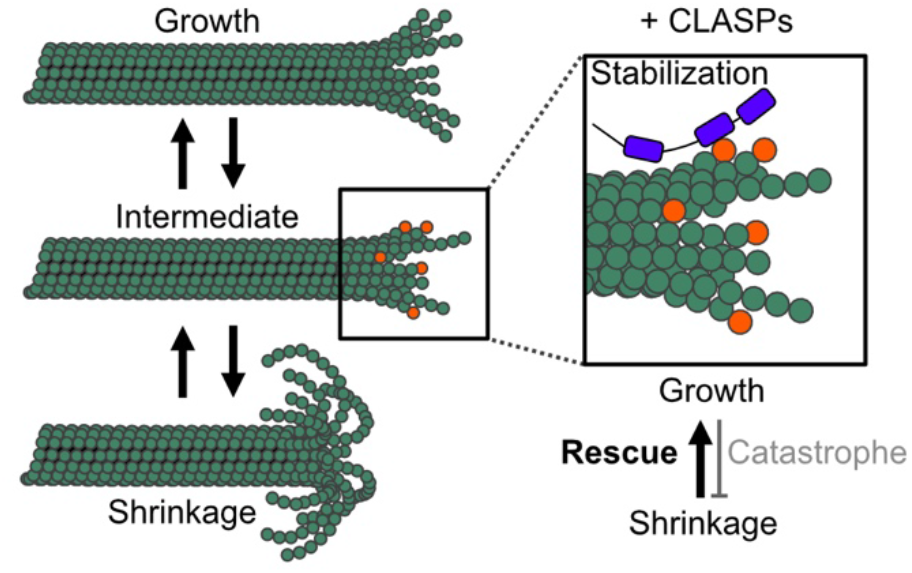
CLASPs stabilize an intermediate state between microtubule growth and shrinkage. Microtubules switch between growth and shrinkage through a metastable intermediate state, characterized by a unique nucleotide-dependent configuration of tubulin at the microtubule end (denoted in orange). CLASPs (purple) recognize and stabilize this intermediate state, thus suppressing catastrophe and promoting rescue.

What is the nature of the microtubule end in the intermediate state between growth and catastrophe? Our finding that CLASP promotes slow microtubule depolymerization in the presence of GTP, GTPγS, and GDP, but not GMPCPP, suggests that the intermediate state requires a post-GTP hydrolysis state. Indeed, it has been proposed that the transient exposure of terminal GDP-tubulin dimers during growth serves as a precursor to catastrophe (Bowne-Anderson et al. 2013, Farmer and Zanic 2022). Exposure of GDP-tubulin at growing microtubule ends has recently been linked to an increase in microtubule growth fluctuations and slowed microtubule growth (Cleary et al. 2022). Furthermore, exchanging GDP for GTP on the microtubule end was predicted to suppress microtubule catastrophe (Piedra et al. 2016). Notably, CLASP does not localize to shrinking microtubule ends, which depolymerize through the loss of GDP-tubulin subunits. Therefore, CLASP likely recognizes a distinct microtubule end configuration specific to the pre-catastrophe state, which may involve exposed GDP-tubulin subunits. Importantly, we find that the minimal TOG2 construct recapitulates CLASP’s depolymerase activity on stable microtubules. Given that TOG2 has a unique architecture that permits binding to a distinct, highly curved conformation of tubulin (Leano et al. 2013, Maki et al. 2015, Leano and Slep 2019, Lawrence et al. 2020), we speculate that this unique tubulin conformation is transiently probed by microtubule ends in the intermediate state (Fedorov et al. 2019). The binding of CLASP could further stabilize this curved tubulin conformation, potentially facilitating nucleotide exchange in tubulin dimers at the microtubule end.

Our data using different nucleotides and analogues indicate that CLASP drives the microtubule into and out of the intermediate state in a nucleotide-dependent manner. Importantly, we found that GMPCPP, a slowly hydrolyzable analog of GTP, does not allow CLASP1-mediated microtubule depolymerization in any of the conditions tested. This result is particularly striking in the tubulin dilution experiments with dynamic microtubules where control microtubules undergo catastrophe within seconds following dilution, but are completely stabilized with CLASP1 and GMPCPP. In the three-state model of microtubule dynamics (Tran et al. 1997), we interpret this as a return of the microtubule end to a fully GTP-like state compatible with growth. Because GMPCPP on its own is not sufficient to prevent dilution-induced microtubule catastrophe, this result gives support to the model in which CLASP1 directly facilitates nucleotide exchange at the microtubule end.

Notably, a recent study reconstituting CLASP2-mediated kinetochore attachments reported nucleotide sensitivity of CLASP-microtubule attachments (Luo et al. 2022). Here the authors used DNA origami to design clusters of CLASP2, which formed sustained load-bearing attachments to stable microtubule ends. The authors found that CLASP’s attachment to microtubule ends was abrogated in the presence of GTP. This finding is consistent with our results suggesting that CLASP recognizes nucleotide-specific microtubule end configuration. Given that CLASPs are critical for stabilizing microtubule ends at kinetochores (Maiato et al. 2002, Maiato et al. 2005, Mimori-Kiyosue et al. 2006, Kolenda et al. 2018), CLASPs may prevent kinetochore-anchored microtubules from undergoing catastrophe by stabilizing the intermediate state. While we speculate that nucleotide exchange in the terminal tubulin dimers likely underlies CLASP’s activity, we cannot rule out a possibility that the conformation of CLASP itself is modulated by GTP. Along these lines, a previous study identified putative GTP-binding motifs in Orbit, the Drosophila homologue of CLASP (Inoue et al. 2000). However, these motifs are only partially conserved in human CLASP proteins, and mutating the remaining conserved residues in human CLASP2 does not impact CLASP2’s activity (Luo et al. 2022). Future studies will be required to distinguish between these possibilities and dissect the underlying mechanisms.

Taken together, our results support a mechanism whereby CLASPs stabilize a nucleotide-dependent intermediate state of the microtubule end, which occurs as the microtubule transitions from growth to catastrophe. For dynamically growing microtubules, extending the period of time spent in the intermediate state would allow the microtubule to re-enter the growth phase, thus avoiding catastrophe. We further propose that stabilizing the intermediate state underlies CLASP’s mechanism of microtubule rescue. For example, lattice defects could transiently stabilize the fast-shrinking ends after catastrophe, prompting a return to the intermediate state, which is recognized and stabilized by CLASP. Indeed, previous work showed that CLASP promotes lattice repair and that just a few CLASP molecules are sufficient to promote microtubule rescue (Aher et al. 2018, Aher et al. 2020). We, therefore, present a unifying mechanism underlying the two major activities of CLASP in suppressing catastrophe and promoting rescue. Overall, our discovery that CLASP1 stabilizes the intermediate state between microtubule growth and shrinkage in a nucleotide-dependent manner provides key mechanistic insights into an important family of microtubule regulatory proteins. To what extent other microtubule-associated proteins regulate the stability of the intermediate state is an exciting area for future research.

## METHODS

### DNA constructs

Human His-CLASP1 (NM_015282.2) in pFastBacHT vector was purchased from Genscript. The cDNA encoding full-length human CLASP2α was purchased from Dharmacon (Accession: BC140778.1) and subcloned into a pFastBacHT vector (Invitrogen) containing an N-terminal 6xHis-tag. The His-CLASP2γ construct was generated as previously described (Lawrence et al., 2018). The cDNA encoding His-EGFP-L-TOG2-S in a pRSETa vector was a gift from E. Grishchuk (University of Pennsylvania, Philadelphia, PA, USA) (Luo et al. 2022). The plasmid containing chTOG cDNA was a gift from Stephen Royle (Addgene plasmid # 69108; http://n2t.net/addgene:69108; RRID: Addgene_69108). The chTOG cDNA was subcloned into a modified pFastBac vector containing an N-terminal 6xHis tag (a gift from G. Brouhard, McGill University, Montréal, QC, Canada). Cloning products were verified by DNA sequencing.

### Protein preparation

#### Tubulin purification and fluorescent labeling

Bovine brain tubulin was purified using cycles of polymerization and depolymerization using the high-molarity PIPES method (Castoldi and Popov 2003). Tubulin was labeled with tetramethylrhodamine (TAMRA) and Alexa Fluor 488 dyes (ThermoFisher Scientific, Waltham, MA, USA) according to the standard protocols and as previously described (Hyman et al. 1991, Gell et al. 2010). Fluorescently-labeled tubulin was used at a ratio of between 5% and 25% of the total tubulin.

#### CLASP and chTOG protein purification

His-CLASP1 protein was expressed in baculovirus-infected Sf9 insect cells using the Bac-to-Bac system (Invitrogen). After the first amplification, baculovirus-infected insect cells (BIIC) stocks were used to infect Sf9 cells at a density of 1 × 10^6^ viable cells/ml at a ratio of 10^−4^ BIIC:total culture volume (Wasilko and Lee 2006, Wasilko et al. 2009). Cells were harvested 5 days after infection. His-CLASP1 cell pellets were lysed by one freeze–thaw cycle and Dounce homogenizing in lysis buffer containing 50 mM HEPES (4-(2-hydroxyethyl)-1-piperazineethanesulfonic acid) (pH 7.5), 150 mM NaCl, 2 mM MgCl_2_, 5% (v/v) glycerol, 0.1% (v/v) Tween-20, 2 mM MgCl_2_, 10 mM imidazole, 1 mM dithiothreitol (DTT) and supplemented with protease inhibitors. Genomic DNA was sheared by passing the lysate through an 18-gauge needle. The crude lysates were clarified by centrifugation for 20 min at 4°C and 35,000 rpm in a Beckman L90K Optima and 50.2 Ti rotor. Clarified lysates were applied to a HisTrapHP column (GE Lifesciences) according to the manufacturer’s protocol. His-CLASP1 protein was eluted in 50 mM HEPES (pH 6.8), 150 mM NaCl, 5% (v/v) glycerol, 0.1% (v/v) Tween-20, 2 mM MgCl_2_, 1 mM DTT, 50 mM L-glutamate, 50 mM L-arginine, and a linear gradient of 50 mM - 300 mM imidazole. His-CLASP1 was further purified and buffer exchanged into CLASP storage buffer (25 mM PIPES [piperazine-N,N′-bis(2-ethanesulfonic acid)] [pH 6.8], 150 mM KCl, 5% (v/v) glycerol, 0.1% (v/v) Tween-20, 50 mM L-glutamate, 50 mM L-arginine, and 1 mM DTT) by size exclusion chromatography using a Superdex 200 Increase 10/300 GL column (Cytiva). Purified CLASP1 was labeled using Alexa Fluor 488 Microscale Protein Labeling Kit (ThermoFisher Scientific, cat. #A30006) according to the manufacturer’s instructions.

His-CLASP2α protein was expressed in Sf9 insect cells and purified as described above for His-CLASP1 with the following modifications. Cell pellets were lysed in 50 mM PIPES (pH 6.8), 120 mM KCl, 2 mM MgCl_2_, 50 mM L-glutamate, 50 mM L-arginine, 10% glycerol (v/v), 0.1% (v/v) Tween-20, 1 mM DTT and supplemented with protease inhibitors. His-CLASP2α was eluted in 50 mM PIPES (pH 6.8), 400 mM KCl, 5% (v/v) glycerol, 0.1% (v/v) Tween-20, 2 mM MgCl_2_, 1 mM DTT, 50 mM L-glutamate, 50 mM L-arginine, and a linear gradient of 50 mM - 300 mM imidazole. Purified His-CLASP2α from the HisTrap column was desalted into CLASP storage buffer using an Amicon centrifugal filter. His-CLASP2γ protein was expressed in Sf9 insect cells and purified as previously described (Lawrence et al. 2018).

His-EGFP-L-TOG2-S protein was expressed in BL21(DE3) E. *coli* cells. Expression was induced with 0.2 mM IPTG at 18°C for 16 h. Cells were lysed for 1 hr at 4°C in 50 mM HEPES (pH 7.5), 300 mM NaCl, 2 mM MgCl_2_, 5% (v/v) glycerol, 0.1% (v/v) Tween-20, 1 mM DTT and 40 mM imidazole and supplemented with 1 mg/ml lysozyme, 10 mg/ml PMSF and EDTA-free protease inhibitors (Roche). The crude lysate was sonicated on ice and then clarified by centrifugation for 30 min at 4°C and 35,000 rpm in a Beckman L90K Optima and 50.2 Ti rotor. Clarified lysates were applied to a HisTrapHP column (Cytiva) according to the manufacturer’s protocol. His-EGFP-L-TOG2-S protein was eluted with 50 mM HEPES (pH 7.5), 500 mM NaCl, 2 mM MgCl_2_, 5% (v/v) glycerol, 0.1% (v/v) Tween-20 and 1 mM DTT and linear gradient of 40 mM - 500 mM imidazole. Purified His-EGFP-L-TOG2-S from the HisTrap column was desalted into CLASP storage buffer using a PD-10 desalting column (Cytiva).

His-chTOG protein was expressed in Sf9 insect cells and purified as described above for His-CLASP1 with the following modifications. Cell pellets were lysed in buffer containing 50 mM HEPES (pH 7.2), 400 mM KCl, 2 mM MgCl_2_, 10% glycerol, 0.005% Brij35, 1 mM dithiothreitol (DTT) and supplemented with protease inhibitors. His-chTOG protein was eluted in 50 mM HEPES (pH 7.2), 400 mM KCl, 2 mM MgCl_2_, 10% (v/v) glycerol, 0.005% (v/v) Brij35, 1 mM dithiothreitol (DTT), and a linear gradient of 60 mM – 400 mM imidazole. His-chTOG was further purified and buffer exchanged into storage buffer (50 mM HEPES [pH 7.2], 400 mM KCl, 2 mM MgCl_2_, 10% (v/v) glycerol, 0.005% (v/v) Brij35, 1 mM DTT) by size exclusion chromatography using a Superdex 200 Increase 10/300 GL column (Cytiva).

Protein purity was assessed by SDS-PAGE and/or mass spectrometry analysis. All proteins were snap frozen in liquid nitrogen as single-use aliquots and stored at −80°C.

### TIRF microscopy and microfluidic channel preparation

Imaging was performed using a Nikon Eclipse Ti microscope with a 100×/1.49 n.a. TIRF objective (Nikon, Tokyo, Japan), Andor iXon Ultra EM-CCD (electron multiplying charge-coupled device) camera (Andor, Belfast, UK); 488- and 561-solid-state lasers (Nikon Lu-NA); HS-625 high-speed emission filter wheel (Finger Lakes Instrumentation, Lima, NY USA); and standard filter sets. For the data in Figure S2, images were acquired with a Nikon high speed Ti-E microscope with epifluorescence, a 100x/1.49 TIRFApo objective and an Andor Neo 5.5 sCMOS camera (2560×2160 pixels, 1 pixel = 6.5 μm x 6.5 μm). In both cases, the microscopes were computer-controlled using Nikon Elements. A Tokai-Hit objective heater was used to maintain the sample at 35°C for all experiments. Images were acquired with exposure times of 50 ms – 200 ms and at the frame rates specified in the methods.

Microscope chambers were constructed as previously described (Gell et al. 2010). Briefly, 22 × 22 mm and 18 × 18 mm silanized coverslips were separated by strips of Parafilm to create narrow channels for the exchange of solution. The channels were rinsed with BRB80 (80 mM PIPES adjusted to pH 6.8 with KOH, 1 mM MgCl_2_, and 1 mM EGTA), incubated with 1:50 anti-TRITC antibody (ThermoFisher Scientific. #A-6397) in BRB80 for 5 min, rinsed with BRB80, incubated with 1% Pluronic F127 in BRB80 for 30 min and rinsed again with BRB80.

### GMPCPP-stabilized microtubule depolymerization assay

GMPCPP-stabilized microtubules (TAMRA-labeled) were prepared according to standard protocols (Gell et al. 2010). Microtubules were introduced into the imaging chamber and adhered to anti-TRITC antibody-coated coverslip surfaces. Reaction mixes containing concentrations of the proteins and nucleotides specified in the main text were introduced into the channels in imaging buffer. The imaging buffer consisted of BRB80 supplemented with 40 mM glucose, 40 μg/ml glucose oxidase, 16 μg/ml catalase, 0.5 mg/ml casein, 50 mM KCl, 10 mM DTT and 1 mM MgCl_2_. Microtubules were imaged for 15 minutes at a frame rate of 0.04 fps. Microtubule lengths were tracked over time with FIESTA (Ruhnow et al. 2011) and used to calculate mean depolymerization rates for the first 5 minutes of depolymerization. For each individual microtubule, the filament length in every frame for the first 5 minutes of the movie (0.04 fps) was plotted against time in MATLAB, and the depolymerization rates were determined from the slope of the linear regression line fitted to the data. Outliers were identified as ± 3 x SD away from the mean and discarded. Data were plotted as “SuperPlots” in which individual data points are color-coded by experimental repeat (Lord et al. 2020). Statistical significance testing was performed on the means of the experimental repeats using a paired two-tailed t-test, or one-way ANOVA on the pooled data followed by a post hoc Tukey HSD as specified in the text.

### Polarity-marked microtubule depolymerization assay

Polarity-marked microtubules were generated by polymerizing A488-labeled GMPCPP-tubulin extensions from TAMRA-labeled microtubule seeds with 3.5 μM - 5 μM A488-tubulin and 1 mM GMPCPP for 10 minutes in microfluidic channels. Since microtubules grown with GMPCPP do not undergo catastrophe and microtubule plus ends grow faster than minus ends, the two ends were distinguished by their lengths, with the longer extension designated as the plus end and the shorter extension as the minus end. Reaction mixes containing CLASP1 and nucleotides as indicated in the main text were introduced into the channels in imaging buffer, and microtubule depolymerization was imaged for 20 minutes at a frame rate of 0.04 fps. Microtubule depolymerization rates were determined independently for each end of the A448-GMPCPP-stabilized extensions on kymographs over the first 5 minutes of the experiment. Outliers were identified as ± 3 x SD away from the mean and discarded. The depolymerization rates of plus and minus ends for the titrations of CLASP1, GTP, and GDP (plus-end only) were fit to the Michaelis-Menten equation:

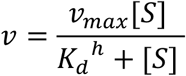

where *v*_*max*_ is the maximum depolymerization rate, *K*_*m*_ is the concentration at which half-maximum rate is achieved

### Tubulin dilution experiments

Microtubules with mixed-nucleotide and GDP lattices were polymerized in the microfluidic channels from GMPCPP-stabilized microtubule seeds. Mixed lattice extensions were grown with 7 μM A488-tubulin and a nucleotide mixture containing 0.2 mM GTP and 0.8 mM GMPCPP for 20 minutes, resulting in microtubule lattices with an estimated nucleotide content of approximately 50% GTP and 50% GMPCPP (Tropini et al. 2012). GDP microtubules were grown with 12 μM A488-tubulin and 1 mM GTP for 10 minutes. Buffer exchange was performed using filter paper to dilute soluble tubulin and introduce reaction mixtures containing nucleotides with or without 23 nM CLASP1 while imaging. Post-dilution, mixed-lattice microtubules were imaged for 15 minutes at a frame rate of 0.2 fps, and GDP-lattice microtubules were imaged at a frame rate of 1 fps for 5 minutes. Microtubule plus-end depolymerization rates were determined from kymographs for the first 5 minutes after dilution. Outliers were identified as ± 3 x SD away from the mean and discarded. Data were plotted as “SuperPlots” as described above and statistical significance testing was performed on the means of the experimental repeats using a paired two-tail t-test.

### Single-molecule dwell time analysis

GMPCPP-stabilized microtubules were incubated for 2 minutes with 200 pM EGFP-L-TOG2-S or 1 nM A488-labelled CLASP1 to allow the reaction to equilibrate in the imaging chambers prior to imaging. For, EGFP-L-TOG2-S, images were acquired at 20 fps for 30 seconds using the maximum 488 nm laser power and 50 ms exposure. For 488-CLASP1, images were acquired at 2 fps for 5 minutes. An image of the microtubule seed was taken before and after the timelapse. The durations of the binding events on microtubule ends were measured from kymographs generated in Fiji (Schindelin et al. 2012) by manually marking the beginning and end of each event. Dwell times were plotted as histograms with 0.2 s (4-frame) bins and cumulative distribution frequency (CDF) plots in MATLAB. Statistical significance was determined using a Mann-Whitney test.

### Computational simulations

To understand how the process of CLASP-mediated nucleotide-dependent depolymerization progresses over time, we traced the experimental kymographs of polarity-marked GMPCPP-stabilized microtubules incubated with 61.5 nM CLASP and 1 mM GTP and looked at the time dependence of depolymerization rates at microtubules minus and plus ends (Figure S2).

We modeled microtubule depolymerization by incorporating the feature of nucleotide exchange at tubulin dimers incorporated in a microtubule. To implement the exchange kinetics at tubulin dimer level in the model, we first re-shape/project the hollow cylindrical manifold of a microtubule onto a 2-dimensional lattice where each lattice node represents a site for the tubulin dimer (Figure S1A). Of note, the 2-dimensional lattice is a rectangular domain of size *L*_*x*_ *X L*_*y*_ where *L*_*x*_ denotes the number of tubulin dimers in a single protofilament, and *L*_*y*_ denotes the total number of protofilaments (*N*_*pf*_) in the microtubule under consideration (typically 14 for GMPCPP-stabilized microtubules, see Figure S1A). For simplicity, we considered that all the protofilaments in a microtubule are of same length as the microtubule itself. Each lattice site can accommodate at most a single tubulin dimer. The nucleotide exchange at a site (either at microtubule lattice or at microtubule ends or both) was considered as a ‘change of state’ of a tubulin dimer residing at the site under consideration (as illustrated in Figure S1A, where upon nucleotide exchange, ‘green’ sites turn into ‘blue’).

In simulations, the nucleotide exchange happens with pre-defined rates at tubulin dimers incorporated at the microtubule ends as well as at the tubulin dimers embodied within the microtubule lattice. In experiments, we observed that in presence of a fixed concentration of GTP, the depolymerization rate at microtubules ends increases with increasing CLASP concentration (Figure 3). This observation led us to consider the rate of exchange in simulations to be proportional to the CLASP concentration. However, in experiments, in absence of CLASP (i.e zero CLASP concentration), the ends also depolymerize, albeit at a much slower rate. Henceforth, there would be another concentration-independent timescale associated with the exchange kinetics. Taken together, the effective exchange rate at plus/minus end was considered to be 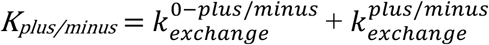 [M] where [M] is the effective CLASP concentration and 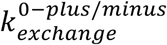 is the CLASP concentration independent nucleotide exchange rates at microtubule ends. Further, the probability of exchange 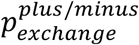 is defined as (1 – exp(−*K*_*plus/minus*_ Δt)) where Δt denotes the time step chosen for the simulation.

In each iteration of the simulation, we scanned the columns at the microtubule ends and randomly visited all *N*_*pf*_ number of sites along the rows. Upon visiting a randomly picked microtubule end site, we drew a random number *p* between 0 and 1 from uniform distribution. If *p* is less than 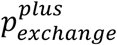 at the plus end (or 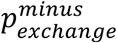 at the minus end), we considered that the nucleotide exchange had occurred at the site under consideration. Further we considered that a terminally exposed column would fall off from a microtubule end if a critical number of tubulin dimers are exchanged at the exposed column. Through this iterative exchange process, if the number of exchanged sites at an exposed end column equals/exceeds a predetermined threshold value (*N*_*threshold*_), that end column falls off. For our simulations, if not mentioned otherwise, *N*_*threshold*_ was chosen to be 6 (Atherton et al. 2018). In the model, removal of a terminally exposed end column marks the depolymerization of a linear array of tubulin dimers at microtubule end which shortens the microtubule length by 8 nm-the length of a single tubulin dimer (Figure S1A).

Next, we introduced the nucleotide exchange at tubulin dimers integrated within microtubule lattice at a pre-defined rate in addition to the exchanges that are occurring at the microtubule ends. The rationale underlying this consideration is described in the following. At plus ends, CLASP may facilitate nucleotide exchange from GMPCPP to GTP which acts as a precursor for terminal tubulin removal and depolymerization. However, at minus ends, the exchangeable nucleotide in the β-tubulin subunit is ‘buried’ inside the lattice. Therefore, if the exchange happens at the minus ends, it may also happen within the microtubule lattice.

For the argument posed above, we chose the minus end exchange rate to be the same as the exchange rate on the lattice. The probability of lattice exchange 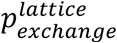 is defined as 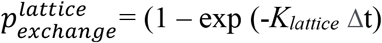 where *K*_*lattice*_ is *k*^lattice^ [M] and *k*^lattice^ denotes the rate of lattice exchange per unit time per unit concentration. For simplicity, in simulations having lattice exchange switched ‘on’, we chose 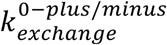 – the CLASP concentration-independent end exchange rates to be zero as its effect was small compared to concentration-dependent end exchange rates. In presence of nucleotide exchange at lattice incorporated tubulin dimers, at a particular time step, if a terminally exposed column of dimers falls off, iteratively the adjacent column (the newly exposed one) also falls off when the number of exchanged sites at that column reaches/exceeds *N*_*threshold*_.

The model parameters are listed in Table S2. The code for the agent-based nucleotide exchange simulations was written in MATLAB (The MathWorks, Natick, MA) and is available at https://github.com/ZanicLab/. A single simulation run of microtubule depolymerization takes minutes of real-time in Apple M1 CPU, RAM 16 GB.

## SUPPLEMENTAL MATERIALS

**Figure S1.**
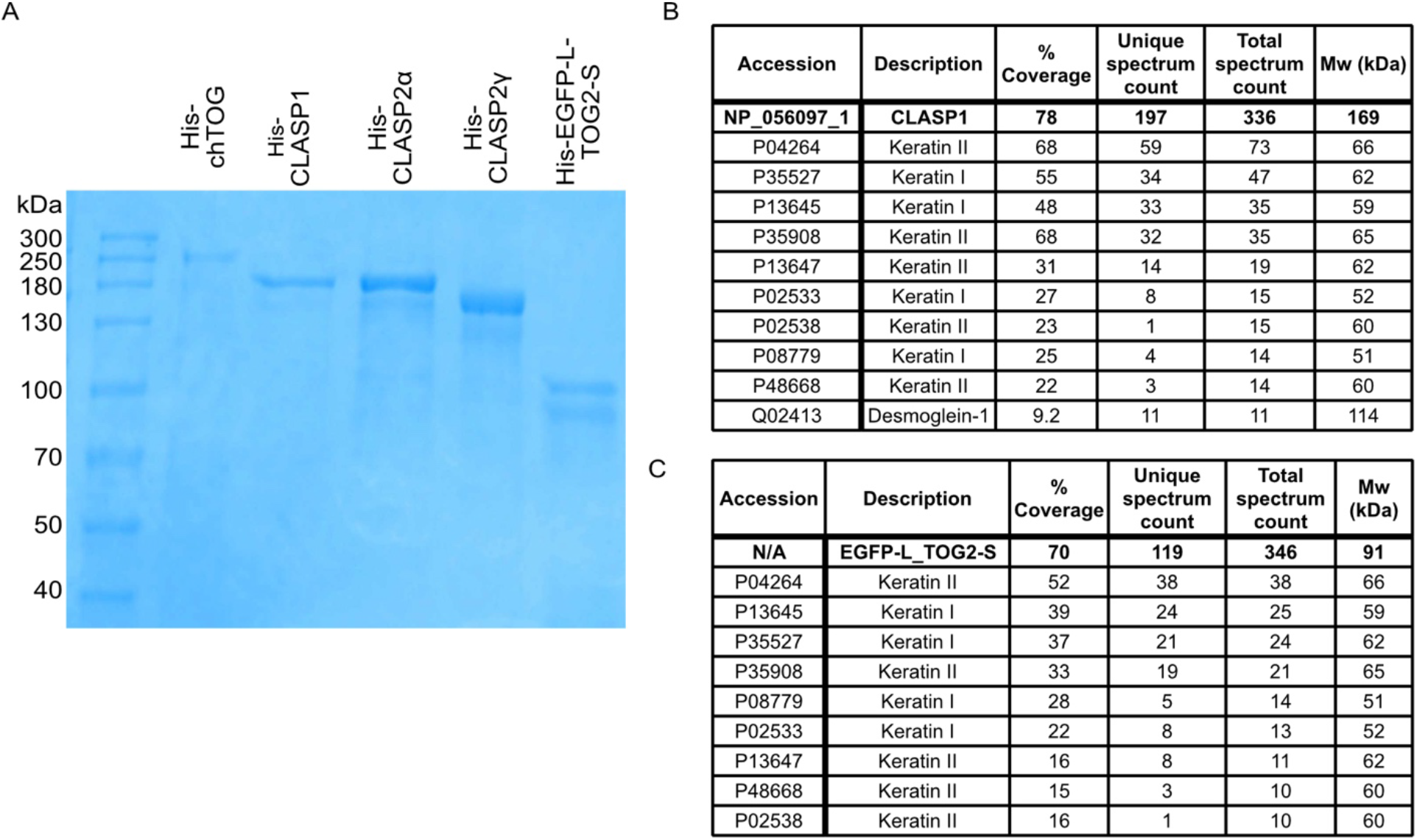
Purification of proteins used in this study. (A) SDS-page gel showing purified His-chTOG, His-CLASP1, His-CLASP2α, His-CLASP2γ and His-EGFP-L-TOG2-S proteins. The lower band in the His-EGFP-L-TOG2-S sample likely represents a truncated protein or breakdown product as no significant contaminating proteins were found in the mass spectrometry analysis. (B) Mass spectrometry analysis of His-CLASP1 and His-EGFP-L-TOG2-S proteins. The hits with a total spectrum count of 10 or more are listed.

**Figure S2.**
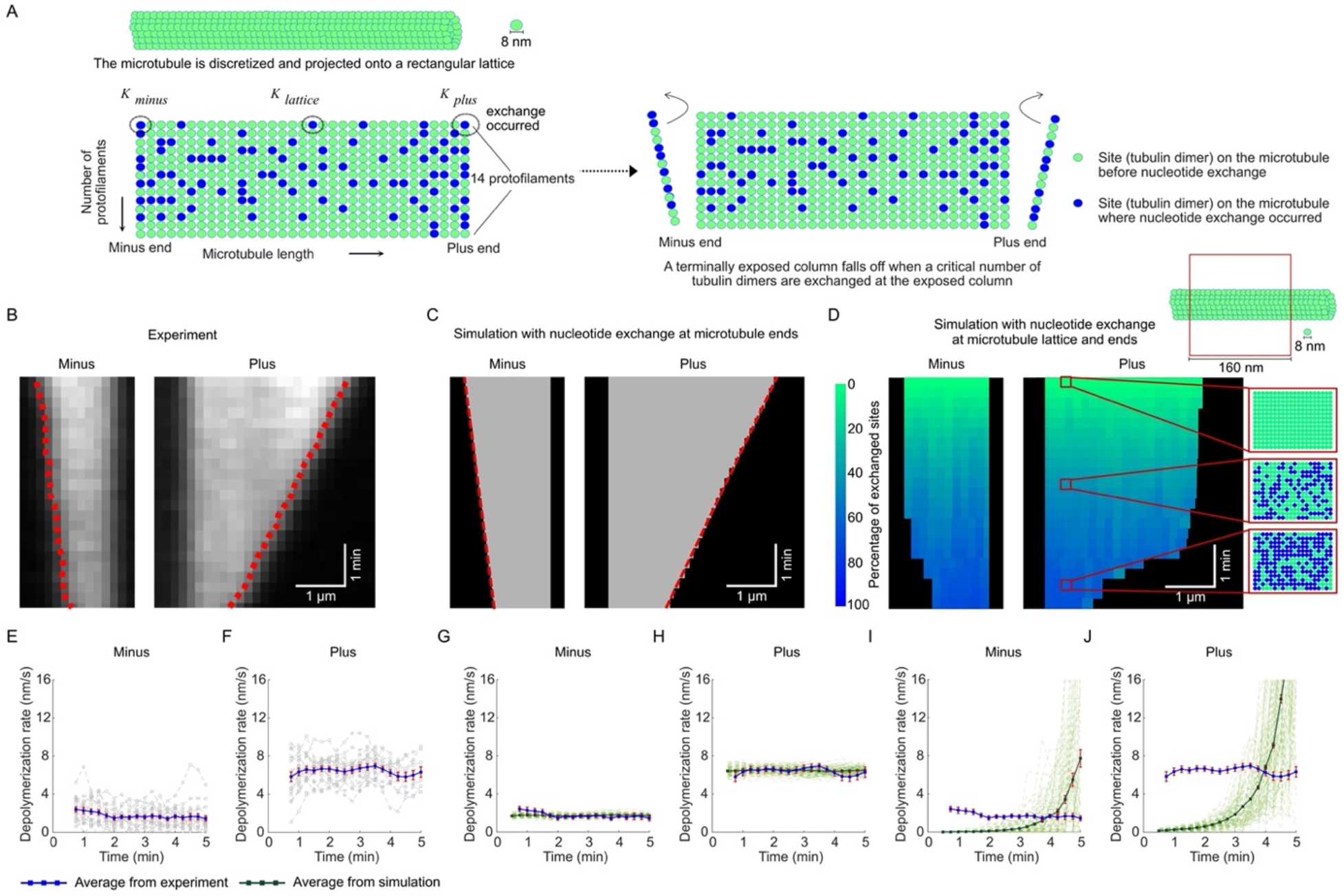
Computational modeling demonstrates that nucleotide exchange in the microtubule lattice does not recapitulate the characteristics of the microtubule depolymerization observed in experiments. (A) Schematics of the computational model for CLASP-dependent nucleotide-exchange leading to depolymerization. (B) A representative kymograph of an experimental microtubule depolymerizing in the presence of 1 mM GTP and 61.5 nM CLASP. The red dashed lines indicate the traces of the microtubule ends obtained using KymographClear and KymographDirect (Mangeol et al., 2016). (C) Representative kymograph shows microtubule depolymerization simulated using the model of nucleotide exchange solely at the microtubule ends. (D) Representative color-coded kymograph shows microtubule depolymerization simulated using model of nucleotide exchange throughout the entire microtubule lattice in addition to the exchange at the ends. The colormap represents the percentage of exchanged sites on the microtubule. The ‘zoomed in’ regions on the kymograph show the tubulin dimers within a 160 nm long segment on the microtubule lattice, at three different time points (0 min, 2.5 min and 5 min). Over the course of time, more and more tubulin dimers are exchanged (‘green’ sites turning into ‘blue’ sites) inside the microtubule lattice. (E-F) The time dependence of depolymerization rate at microtubule minus (E) and plus (F) ends as observed in experiments. In (E-F) the grey curves denote trajectories obtained from individual microtubules in experiment (N=20). The blue curve denotes the average curve evaluated from grey trajectories. (G-H) The time dependence of depolymerization rate at microtubule minus (G) and plus (H) ends as obtained from the model of nucleotide exchange solely at the microtubule ends. (I-J) The time dependence of depolymerization rate at microtubule minus (I) and plus (J) ends as obtained from the model of nucleotide exchange throughout the entire microtubule lattice in addition to the exchange at the ends. In (G-J) the light green curves denote trajectories obtained from individual microtubules in simulation. The dark green curve denotes the average curve evaluated from individual microtubule trajectories in simulation. The instantaneous depolymerization rates presented in (E-J) were estimated by averaging over a 1 min time window. The error bars represent SEM (N=20 in experiment, N=100 in all simulated cases).

**Figure S3.**
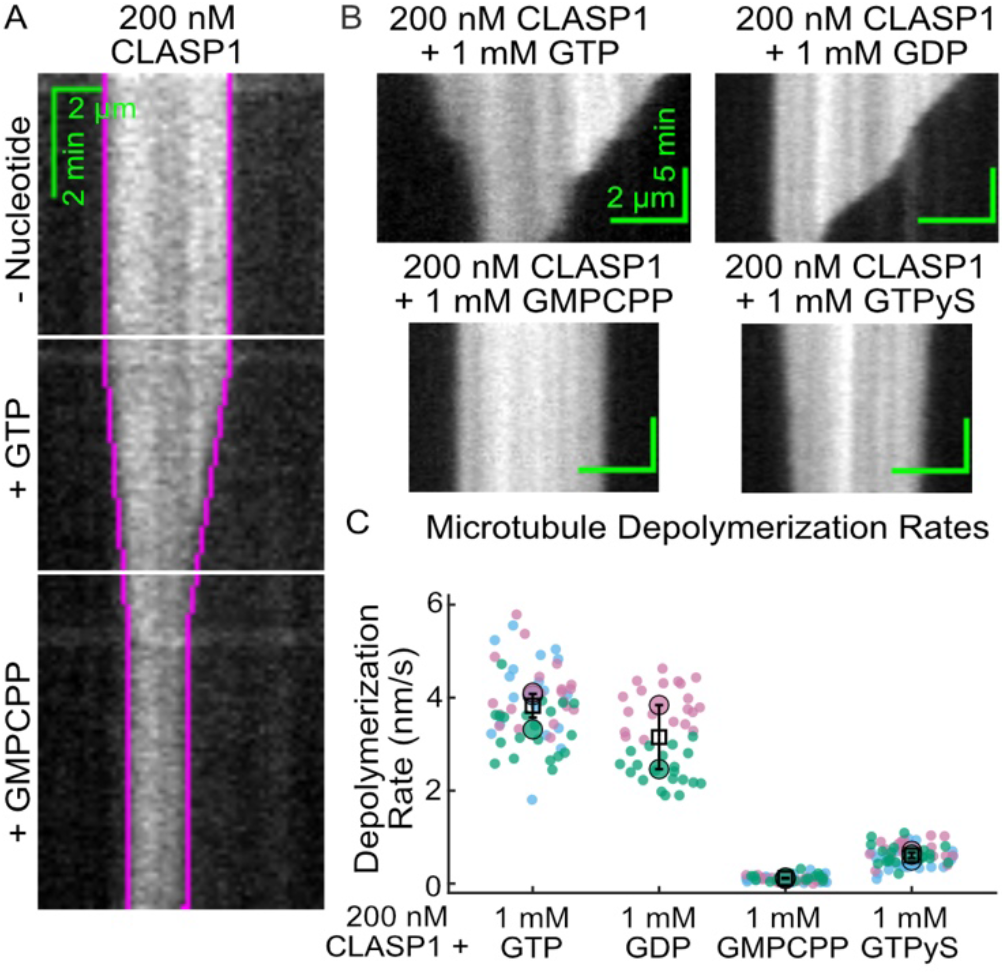
The post-hydrolysis nucleotide state is required for CLASP1’s microtubule depolymerase activity. (A) Representative kymograph of a GMPCPP-stabilized microtubule incubated sequentially with no nucleotide, 1 mM GTP and 1 mM GMPCPP. The white horizontal lines indicate the times of solution exchange; the purple line is a manual trace of the microtubule end position overlaid onto the kymograph. (B) Representative kymographs of GMPCPP-stabilized microtubules incubated with buffer or 200 nM CLASP1 in the presence of 1 mM GDP, 1 mM GMPCPP and 1 mM GTP**γ**S. (C) Quantification of microtubule depolymerization rates for the conditions in the presence of different nucleotides. The mean depolymerization rates were 3.8 nm/s ± 0.3 nm/s; (SE, N=60) with GTP, 3.1 nm/s ± 0.7 nm/s; (SE, N = 40) with GDP, 0.115 +/- 0.008 nm/s; (SE, N=60) with GMPCPP, and 0.61 +/- 0.06 nm/s; (SE, N=60) with GTP**γ**S.

**Figure S4.**
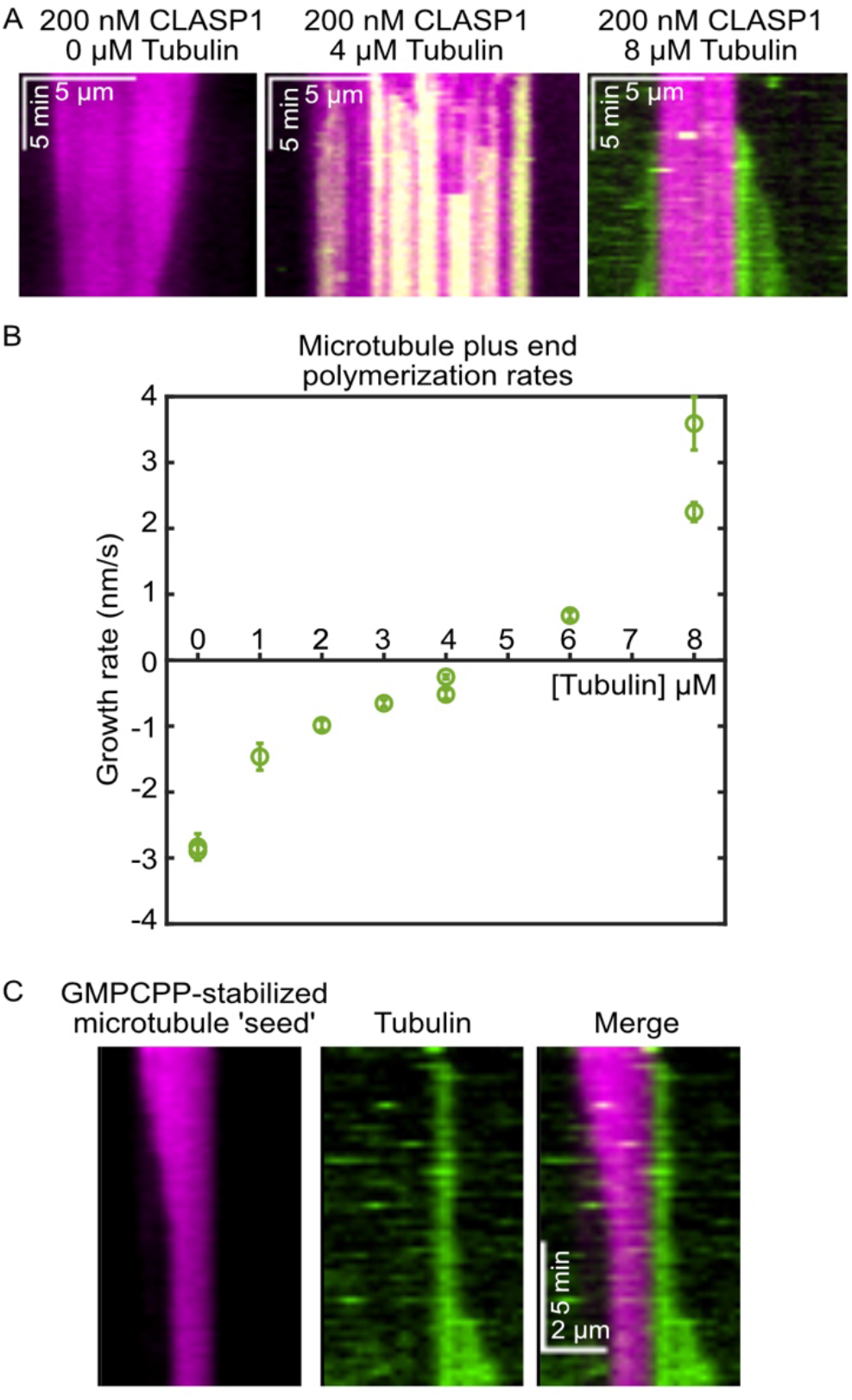
Soluble tubulin does not abolish CLASP’s depolymerase activity until the tubulin concentration is above the critical concentration for microtubule growth. (A) Representative kymographs of microtubules in the presence of 0 μM, 4 μM and 8 μM soluble tubulin and 200 nM CLASP1. The stable microtubule seed is shown in magenta and the tubulin is shown in green. (B) Quantification of the microtubule plus-end polymerization rate across a range of tubulin concentrations from 0 μM to 8 μM in the presence of 200 nM CLASP1. (C) An example kymograph of a microtubule undergoing CLASP-mediated depolymerization at one end polymerization at the other in the presence of 8μM soluble tubulin and 200 nM CLASP1.

**Supplementary Table 1.** Depolymerization rates of different microtubule substrates in the presence and absence of CLASP1. Data related to Figure 5C. Each condition represents data from 3 independent experimental days.

|  | <u>Mixed nucleotide microtubules</u> |  |  |  |  |  |  | <u>GDP microtubules</u> |  |  |  |  |  |  |
| --- | --- | --- | --- | --- | --- | --- | --- | --- | --- | --- | --- | --- | --- | --- |
|  | Control |  |  | CLASP1 |  |  | p value<br>(control<br>versus<br>CLASP1) | Control |  |  | CLASP1 |  |  | p value<br>(control<br>versus<br>CLASP1) |
|  | mean<br>rate<br>(nm/s) | SE | number<br>of MTs | mean<br>rate<br>(nm/s) | SE | number<br>of MTs |  | mean<br>rate<br>(nm/s) | SE | number<br>of MTs | mean<br>rate<br>(nm/s) | SE | number<br>of MTs |  |
| **GTP** | 16 | 2 | 28 | 9.4 | 0.9 | 30 | 0.117 | 115 | 12 | 42 | 11 | 2 | 33 | 0.001 |
| **GDP** | 21 | 4 | 29 | 35 | 6 | 30 | 0.095 | 107 | 19 | 68 | 34 | 2 | 64 | 0.059 |
| **GMPCPP** | 16.8 | 0.7 | 30 | 0.33 | 0.04 | 26 | 0.002 | 92 | 16 | 58 | 1 | 0.3 | 25 | 0.03 |

**Supplementary Table 2.** List of model parameters. Related to Figure S2.

| Parameter | Description | Value | Notes |
| --- | --- | --- | --- |
| $N_{pf}$ | Number of protofilaments | 14 | Matching GMPCPP-microtubules |
| $N_{threshold}$ | Critical number of exchanged dimers within a single column that triggers depolymerization of that column | 6 | Atherton et al., 2018 |
| $\Delta t$ | Simulation time step | 0.01 s | Chosen by numerical experimentation |
| [M] | CLASP concentration | 61.5 nM | Chosen from experimental data |
| $k_{exchange}^{0-plus/minus}$<br>[in the model of nucleotide exchange solely at the microtubule ends] | CLASP concentration independent nucleotide exchange rates at microtubule ends | $0.017 \text{ s}^{-1}$ at plus end; $0.014 \text{ s}^{-1}$ at minus end | Numerically obtained by matching experimental depolymerization rates in absence of CLASP |
| $k_{exchange}^{plus/minus}$<br>[in the model of nucleotide exchange solely at the microtubule ends] | Nucleotide exchange rates at microtubule ends<br>/[time][CLASP concentration] | $0.00678 \text{ s}^{-1}\text{nM}^{-1}$ at plus end,<br>$0.00167 \text{ s}^{-1}\text{nM}^{-1}$ at minus end | Numerically obtained by matching experimental depolymerization rates in presence of 61.5 $\mu\text{M}$ CLASP |
| $k_{lattice}$<br>[in the model of nucleotide exchange throughout the entire microtubule lattice in addition to the exchange at the ends] | Nucleotide exchange rates at microtubule lattice/[time][CLASP concentration] | $0.000056 \text{ s}^{-1}\text{nM}^{-1}$ | Numerically obtained by matching experimental depolymerization rates in presence of 61.5 $\mu\text{M}$ CLASP |
| $k_{exchange}^{plus/minus}$<br>[in the model of nucleotide exchange throughout the entire microtubule lattice in addition to the exchange at the ends] | Nucleotide exchange rates at microtubule ends<br>/[time][CLASP concentration] | $0.00019 \text{ s}^{-1}\text{nM}^{-1}$ at plus end,<br>$0.000056 \text{ s}^{-1}\text{nM}^{-1}$ at minus end | Numerically obtained by matching experimental depolymerization rates in presence of 61.5 $\mu\text{M}$ CLASP |

## SUPPLEMENTAL MOVIE CAPTIONS

**Movie 1**. GMPCPP-stabilized microtubules were incubated with 200 nM CLASP1 in the absence (left) and presence of 1 mM GTP (right).

**Movie 2**. Polarity-marked, GMPCPP-stabilized microtubules were incubated with 1 mM GTP in the absence (left) or presence of 500 nM CLASP1 (right).

## ACKNOWLEDGEMENTS

We thank S. Hall for help with protein purification, A. Maiorov and E. Grishchuk (University of Pennsylvania) for the kind gift of the EGFP-L-TOG2-S expression construct, G. Brouhard (McGill University) for the modified pFastBac expression vector. We also thank H. McDonald and the Vanderbilt Mass Spectrometry Research Center (MSRC) Cores for the mass spectrometry analysis, which was supported in part by Vanderbilt Ingram Cancer Center Resource Share Scholarship 2020-2909607. We thank members of the Zanic lab, A. Olivares, E. Grishchuk and the Vanderbilt Microtubules and Motors Club for discussions and feedback. E.J.L. acknowledges the support of the National Institutes of Health IBSTO training grant T32CA119925. M.Z. acknowledges support from the National Institutes of Health grant R35GM119552.

## Notes

### Competing Interest Statement

The authors have declared no competing interest.

